# VRP Assembler: haplotype-resolved *de novo* assembly of diploid and polyploid genomes using quantum computing

**DOI:** 10.1101/2023.10.19.563028

**Authors:** Yibo Chen, Jun-Han Huang, Yuhui Sun, Yong Zhang, Yuxiang Li, Xun Xu

## Abstract

Precision medicine’s emphasis on individual genetic variants highlights the importance of haplotype-resolved assembly, a computational challenge in bioinformatics given its combinatorial nature. While classical algorithms have made strides in addressing this issue, the potential of quantum computing remains largely untapped. Here, we present the VRP assembler: a novel approach that transforms this task into a vehicle routing problem, a combinatorial optimization problem solvable on a quantum computer. We demonstrate its potential and feasibility through a proof of concept on short synthetic diploid and triploid genomes using a D-Wave quantum annealer. To tackle larger-scale assembly problems, we integrate the VRP assembler with Google’s OR-Tools, achieving a haplotype-resolved assembly across the human major histocompatibility complex (MHC) region. Our results show encouraging performance compared to hifiasm with the phasing accuracy first approaching the theoretical limit, underscoring the promising future of quantum computing in bioinformatics.

## 1. Introduction

Haplotype-resolved *de novo* assembly is computationally hard to obtain and is key to the downstream research such as tracking evolution history, locating individual variants, etc. [1–5]. Workflows of *de novo* assembly at haplotype level typically consist of two stages: phasing and assembly. A phasing process is to group sequencing reads into each haplotype, and an assembly process is conducted in each group to obtain a haplotype-resolved outcome. For diploid genomes like the human genome, a common phasing model is minimum error correction (MEC) [6] with several heuristic algorithms [7, 8]. This model utilizes single nucleotide polymorphism (SNP) to group reads with least conflicts among each other. However, solving a MEC metric is NP-hard, meaning that finding the exact optimal solution for a large scale instance is computationally intractable.

In terms of the assembly process, there are two major classes of algorithms to assemble reads into longer contigs: the overlap-layout-consensus (OLC) paradigm [9] and the de Bruijn graphs (DBG) [10]. These two algorithms convert the assembly problem into two different graph theory problems. The OLC approach is essentially solving a Hamiltonian path problem on an overlap graph, which is NP-complete. On the other hand, the DBG approach aims to find an Eulerian path on a k-mer graph, which is a P problem. Several reviews have conducted comprehensive comparisons between OLC and DBG [11, 12], which suggested that the OLC may perform better with noisy long reads from the third-generation sequencing platform, while the DBG seems to be more suitable with accurate short reads from the next-generation sequencing.

After contig construction, further extending them to longer segments needs to link contigs into scaffolds, via solving a complex linkage graph. The difficulty in linking contigs stems from the interleaving structures [12], in which each contig can have multiple links connecting others, making this problem NP-hard. To achieve a haplotype-resolved assembly for diploid and polyploid genomes, many strategies utilize extra information to enhance the assembly quality, such as integrating pair-end reads [13], trio information [14, 15], ultra-long sequencing reads or Hi-C data [16–18]. These extra information may help clustering contigs by means of cutting links between alleles or enforcing links within the same haplotype, which could lead to another NP problem.

Generally, the haplotype assembly may be CPU and memory consuming, mainly due to the combinatorial nature of the mentioned NP problems. In order to balance accuracy and computational costs, algorithms of haplotype assembly trend to adopt heuristic approaches [15, 18–22]. Some of these algorithms convert the mentioned NP problems into optimization models and obtain efficient performance, such as mapping a phasing process onto a Max-Cut problem [18, 20, 21]. Nevertheless, given the complexity of problems themselves, these algorithms could cost considerably amount of time for a single assembly, even on a high-performance computing cluster.

Recent development in quantum computing shows promising potential to cope with heavy computational tasks[23, 24]. Characteristics of quantum system, like superposition and entanglement of qubits, enable calculations on quantum devices to perform exponential acceleration over classical computing [25]. In the past decade, quantum computing has shown its advantage on specific tasks like Gaussian boson sampling and quantum simulation [26–29]. Other potential applications in various fields, such as industry, finance and life sciences, have also been widely discussed [30–39]. In particular, NP problems in bioinformatics are expected to be solved efficiently by quantum optimization [40].

Quantum optimization can be programmed in various ways, such as quantum annealing [41] or quantum approximate optimization algorithm (QAOA) [42, 43]. Both of them solve optimization problems by sampling the ground state of the equivalent quantum Ising model after analog adiabatic evolution [44], where quantum tunneling effect plays a key role in escaping local optima [41, 44]. Therefore, bioinformatics NP problems could be solved on competent quantum devices, if they can be re-modeled and turned into Ising formulation. For instance, *de novo* assembly problem can be mapped onto the travelling salesperson problem (TSP), in Ising formulation, and programmed on a D-Wave quantum annealer [45–47].

Especially, recent development in simulated quantum annealing of complicated spin-glass dynamics showed a scaling advantage over classical algorithms [23], encouraging us to exploit applications of quantum annealing in bioinformatics.

In this article, we propose a haplotype assembly method to combines both phasing and assembly process into a single optimization model: VRP assembler. The VRP assembler models a haplotype assembly as a vehicle routing problem (VRP), enabling the optimization procedure to be solved on quantum annealers as well as gate-based quantum computers to harness potential quantum acceleration. Given this modeling approach, the VRP assembler is naturally suited for the challenging task of polyploid haplotype-resolved assembly, and it operates effectively under conditions of low sequencing error rate. As a testament to its efficacy, we successfully apply the VRP assembler to reconstruct the human major histocompatibility complex (MHC) region using simulated reads sets of various average lengths. The resulting haplotype-resolved assemblies achieve low switch error rates of around 0.1%, approaching the reachable upper limit of accuracy. Furthermore, these switch error rates are generally lower, as low as 0.03%, than those obtained using hifiasm. The general framework depicted in Figure 1 demonstrates the application of VRP to solve the haplotype assembly problem.

**Fig. 1.**
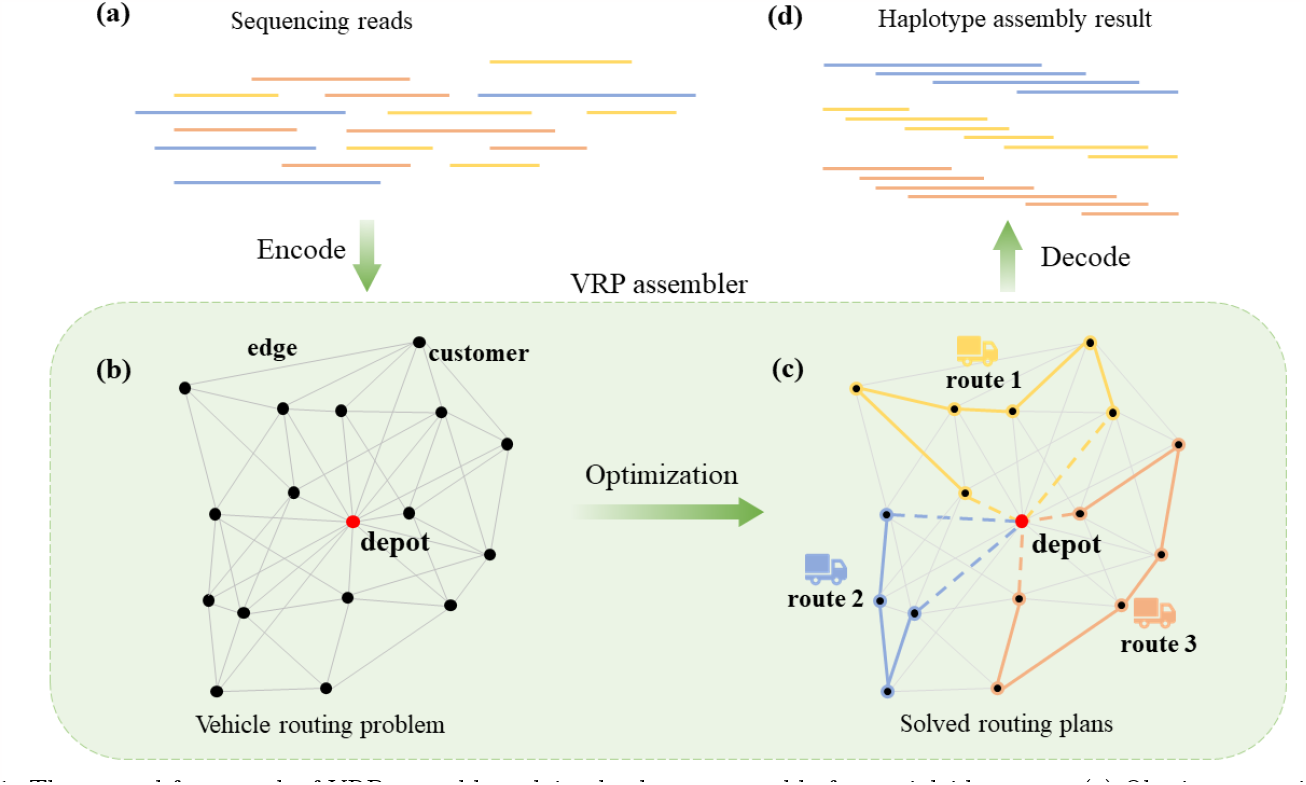
The general framework of VRP assembler solving haplotype assembly for a triploid genome. (a) Obtain sequencing reads. (b) Use pairwise alignment software such as minimap2 to extract overlaps information, and encode reads and overlaps into a VRP model. In this model, reads are seen as customers to visit, while overlaps information are written in the weighted edges. Specifically, the depot is a virtual node and does not represent any real read. (c) Solve VRP instances with optimization solvers. Each vehicle is assigned with several customers to visit in a certain order. In this work, both a D-Wave’s quantum annealing system and Google’s OR-Tools are implemented to solve haplotype assembly in different scales. (d) Decode routing plans from the optimization solver and reconstruct haplotypes according to the assigning and the visiting order of customers. Note that the dashed line connecting the depot should be neglected while decoding since the depot is a virtual read that has no actual alignments with other real reads.

## 2. Results

### 2.1 Overview of VRP assembler

The original VRP is optimal routes planning for a fleet of vehicles delivering goods to customers [48]. The problem is encoded on a graph *G*(*n, E*), with nodes and edges representing the customers and the distance between them, respectively. In VRP assembler, reads are seen as nodes, and the weighted edges are derived from a scoring function of the pairwise alignment information. The VRP assembler relies on the scoring difference brought by heterozygous SNP as well as insertion and deletion (INDEL), so as to distinguish reads from different haplotypes.

As a proof of concept, we perform haplotype assembly tests with short synthetic sequences on a quantum annealer in the D-Wave system. While current quantum annealers are not yet capable of handling the assembly of long sequences with a large number of reads, we have observed encouraging results. Although there are studies suggesting that different ways of embedding and decoding strategies of qubits may enhance the performance of quantum annealer [49, 50], the enhancement still falls shorts of our needs for further experiments. Therefore we leave larger-scale tasks on quantum devices as a future work when more reliable qubits are available. Instead, we implement the VRP assembler with Google’s OR-Tools framework to conduct haplotype assembly on the human major histocompatibility complex (MHC) region. The human MHC region is of ∼5 Mbp long and has dense loci and a high rate of SNP and INDEL, making it a suitable testbed for the VRP assembler. The reference haplotypes are selected from the assembly results reported by Chin *et al*. in Ref. [2], in which the authors achieved a diploid assembly across human MHC region for Genome in a Bottle sample HG002. The results show that the VRP assembler performs admirably, with a lower switch error rate compared to hifiasm’s results. This not only validates the feasibility of our approach, but also implies its potential for improvement over existing heuristic methods as well as the foreseeable utility of quantum computing in bioinformatics.

### 2.2 Haplotype-resolved assembly using quantum annealing

We randomly synthesize short DNA sequences of diploid and triploid genomes and simulate heterozygous SNP. The length of DNA sequences for diploid genomes is set to 50 bp, while that for triploid genomes is 35 bp. We generate reads files from these sequences, with an average length of 20 bp in both cases. Pairwise alignment files in BAM format are generated using Minimap2. These BAM files are then filtered and used to initialize the overlap matrices by the scoring function (Methods). Subsequently, a VRP instance is built from the overlap matrices.

Next, we employ the VRP solver and upload the corresponding QUBO model onto D-Wave’s quantum annealing system. Configuration with lowest energy is retrieved and decoded into the assembly order of artificial reads. The assembly order of reads are further used to reconstruct haplotypes, which are then compared to the original sequences by calculating hamming distance. The entire process are repeated in 10 times for both diploid and triploid genomes. According to the results, the reconstructed sequences exactly match the original sequences with zero hamming distance in all runs, demonstrating the viability of using quantum hardware to solve haplotype assembly problems. Figure 2 shows one of the comparisons between the reconstructed sequences and the original sequences.

**Fig. 2.**
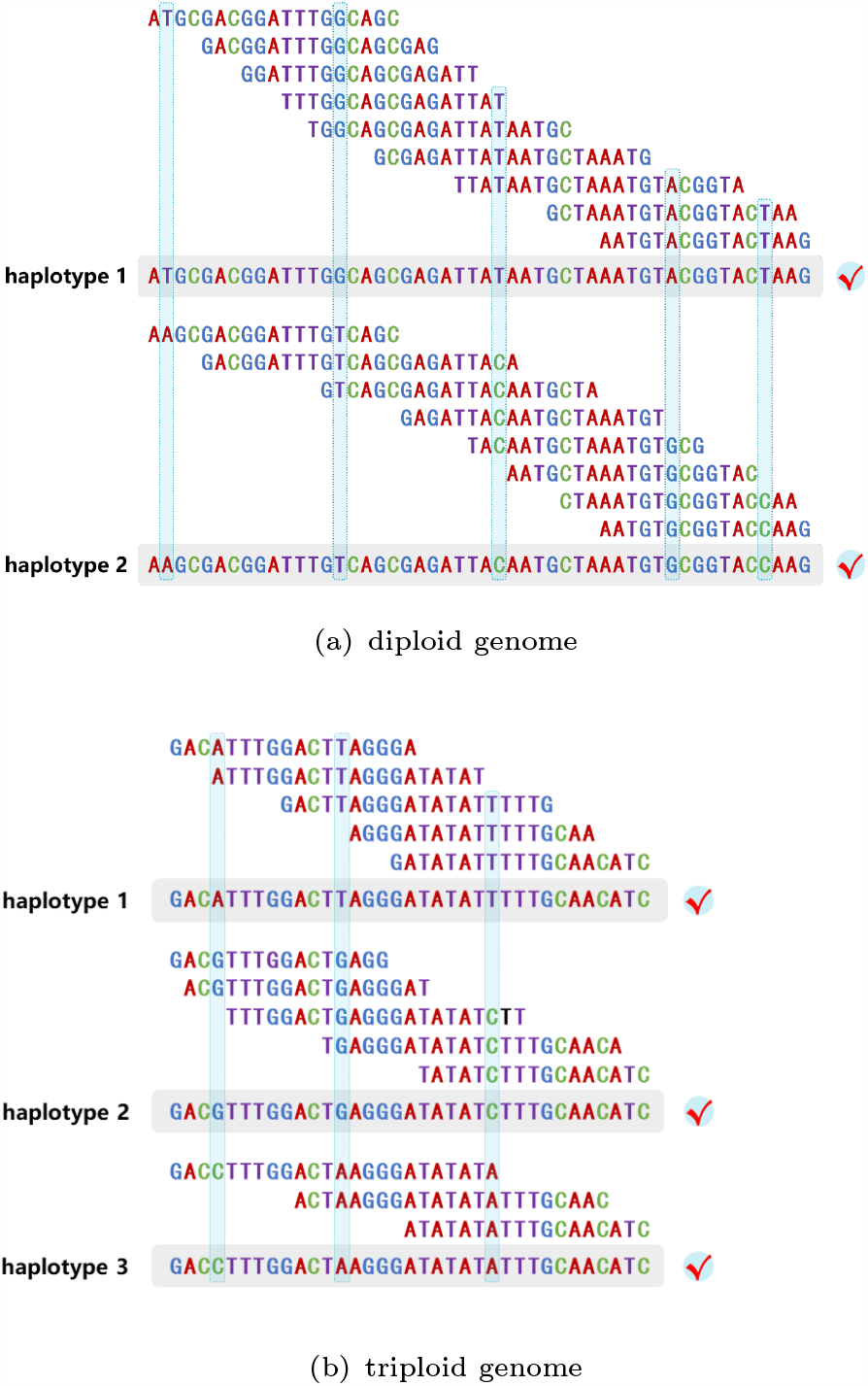
Optimal solution of haplotype assembly in the case of synthetic (a) diploid and (b) triploid sequences. The framed strings are the original sequences. The simulated heterozygous SNP are dash framed. It is easy to see that reconstructed sequences exactly match the original ones.

### 2.3 Benchmarking VRP assembler

The above experiments are conducted without sequencing errors, which could over-estimate the quality of haplotype assembly. However, the scoring function in VRP assembler relies on penalizing mismatches between reads for phasing and assembly, making it naturally error-tolerant by adjusting penalty parameter in our model (Methods), as long as sequencing errors contribute less than heterozygosity. To verify the robustness of the VRP assembler against sequencing errors, we explore the rate of perfect assembly (i.e. all reads are phased accurately and assembled in an expected order, thus no switching between haplotypes) with different sequencing error rate and heterozygosity rate. We use 120 kbp long synthetic DNA sequences of a diploid genome. The reads are generated from synthetic genomes with an average length of 20kbp, close to PacBio HiFi reads. The average length of overlap between reads is set to 12kbp as shown in Fig. 3. This setup exceeds the current capability of the D-Wave quantum annealer with relatively sparse spins connections, prompting us to employ the classical VRP solver in Google’s OR-Tools software. OR-Tools is an open source and integrated framework for solving optimization problems [51], offering flexible searching strategies for customized implementation. In this task, we choose *LocalCheapestInsertion* to generate an initial solution, followed by *GuidedLocalSearch* to escape local minima. The optimization time for each run is set to be 1 second that is enough for the solver to converge.

**Fig. 3.**
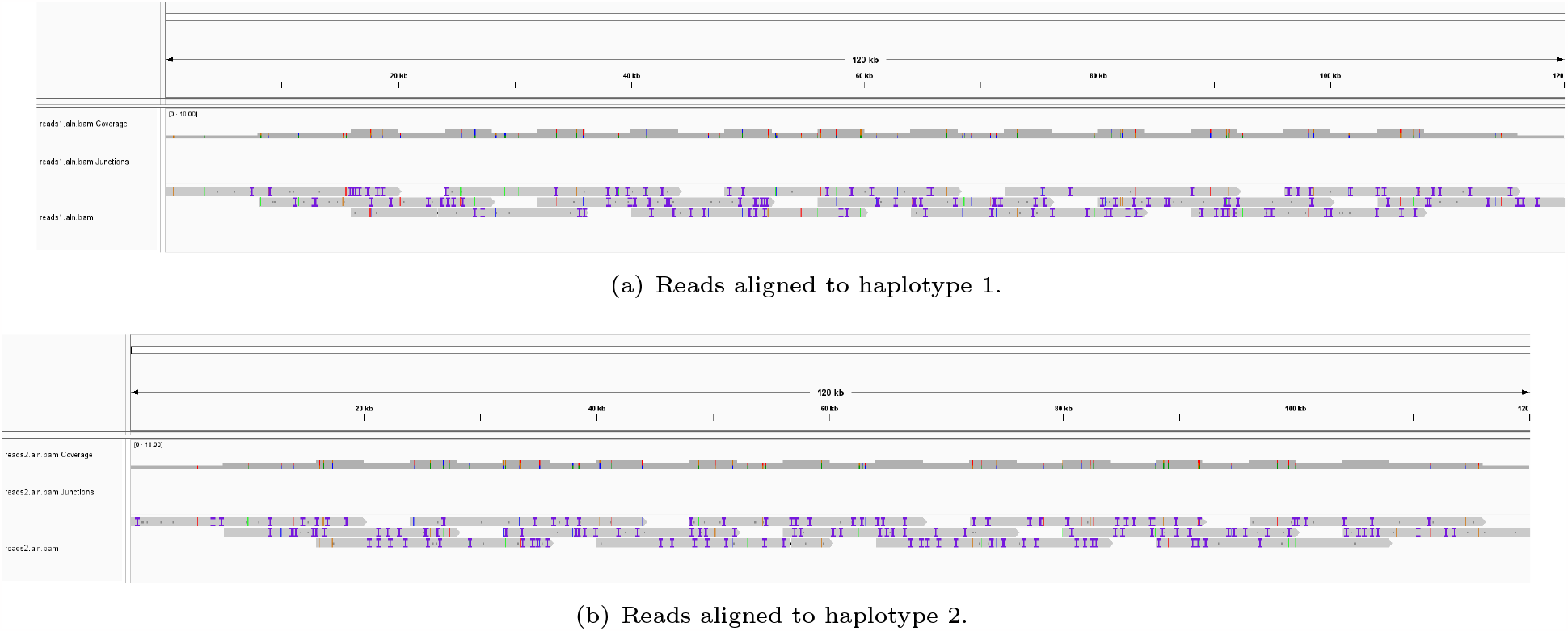
Alignments of generated reads to their original synthetic sequences.

Figure 4 displays our results on a range of different configurations of heterozygosity rate and sequencing error rate. Here, the rate of perfect assembly is defined as the number of times in 100 replicates that the assembly order of the reads exactly matched their order of positions on original sequences. It is straight-forward that a higher heterozygosity rate or a lower error rate will make haplotype assembly easier. Especially, when heterozygosity rate is higher than 0.1%, the VRP assembler can find the perfect assembly in almost every runs, because a higher heterozygosity rate provides more heterozygous sites in the overlapping regions, facilitating the reads connections in the correct order.

**Fig. 4.**
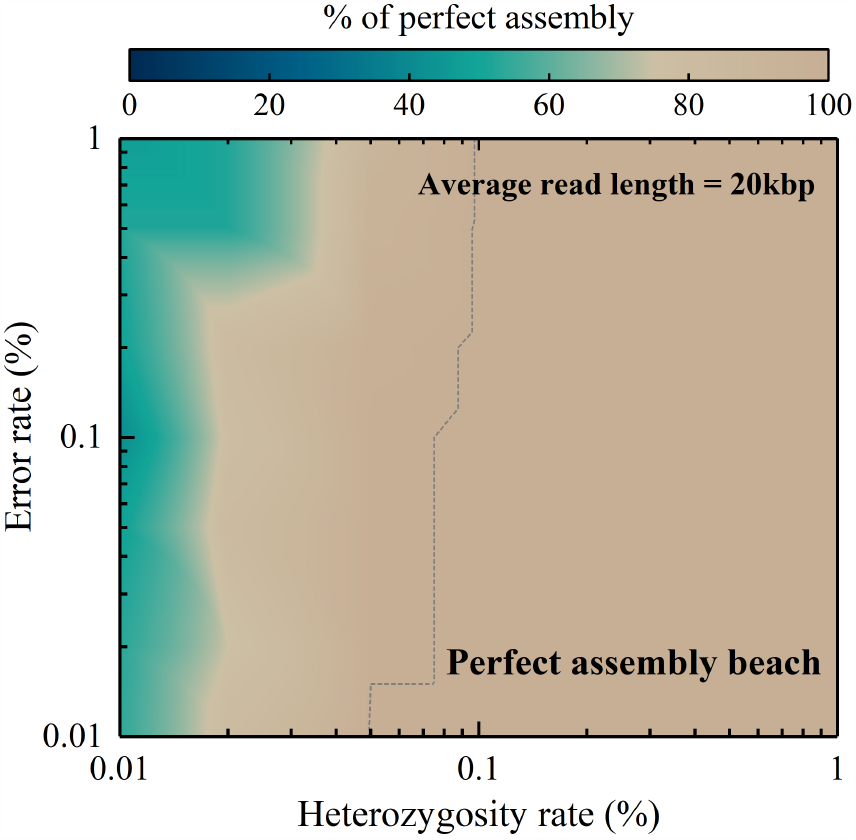
Tests of perfect assembly rate with different heterozygosity rates and sequencing error rates using synthetic reads of 20kbp length. Each data is collected over 100 repetitions. The parameter *p*_1_ and *p*_2_ (Methods) are set to 2 and 0 respectively in all runs. Nevertheless, in actual applications, such as haplotype assembly of real genome, it is necessary to tune both parameters *p*_1_ and *p*_2_, in a concerted manner to ensure optimal outcomes.

### 2.4 Haplotype-resolved assembly of human genomes

The experiments on synthetic DNA sequences only consider heterozygous SNP, whereas real DNA sequences are characterized by a diverse array of variants, including INDELs and repetitive sequences. To further evaluate the performance of VRP assembler on real DNA sequences, we choose reconstruct one of the most SNP-enriched region in the human genome, the MHC region. However, unlike the synthetic genome, the heterozygous sites in MHC region are not uniformly distributed, but rather gathered on several loci. Therefore, it is valuable to test the performance of the VRP assembler on this real DNA region.

The reference haplotype used for reconstruction was sourced from Ref. [2], where the authors used HG002’s 10X linked reads and ONT ultra-long reads to conduct phasing on MHC region. Employing WhatsHap [52] and Peregrine assembler [53], they successfully phased 15k and 20k PacBio HiFi reads together and eventually obtained the haplotype-resolved assembly sequences. From each haplotypes in Ref. [2], we generate three sets of 4 ∼ 6X depth of reads covering the entire MHC region using SimLoRD [54], with 10 simulated reads samples per set. The average read lengths of these three sets are 20kbp, 25kbp, 30kbp, respectively, with a simulated sequencing error of 0.1% [55].

The optimization procedure is performed using OR-Tools, with the total optimization time set to be 300 seconds, but usually the optimal is found before the program ends. The results, consisting of the assembly order of reads for each haplotype, are used to reconstruct a primary assembly of two haplotype sequences. We then use Racon [56] to obtain the consensus sequence as the final output. Since the VRP assembler produce only one contig for each haplotype, resulting in an N50 assembly length equal to the length of the MHC region, as a result, we choose to calculate switch error rate to evaluate the accuracy of phasing. In our implementation, we configure the phasing N50 to the length of MHC region in order to also assess the accuracy of long-range phasing. For comparison, we employ hifiasm [18] to generate de novo assemblies for all three sample sets. The average switch error rate of assemblies from the VRP assembler and hifiasm is calculated per sample and then averaged across all 10 samples in each set.

As depicted in Figure 5, both the VRP assembler and hifiasm perform commendably well in terms of switch error rate. However, as the average read length increases, the switch error rate demonstrates a monotonic decrease for VRP assembler results, while less apparent in hifiasm’s. Besides, the VRP assembler displays marginally superior performance than hifiasm, as evidenced by a consistently lower switch error rate, which, although only 0.01% ~ 0.03% lower than that of hifiasm, is noteworthy. Because the results of the VRP assembler are already approaching the reachable limit, the upper bound of zone II.

**Fig. 5.**
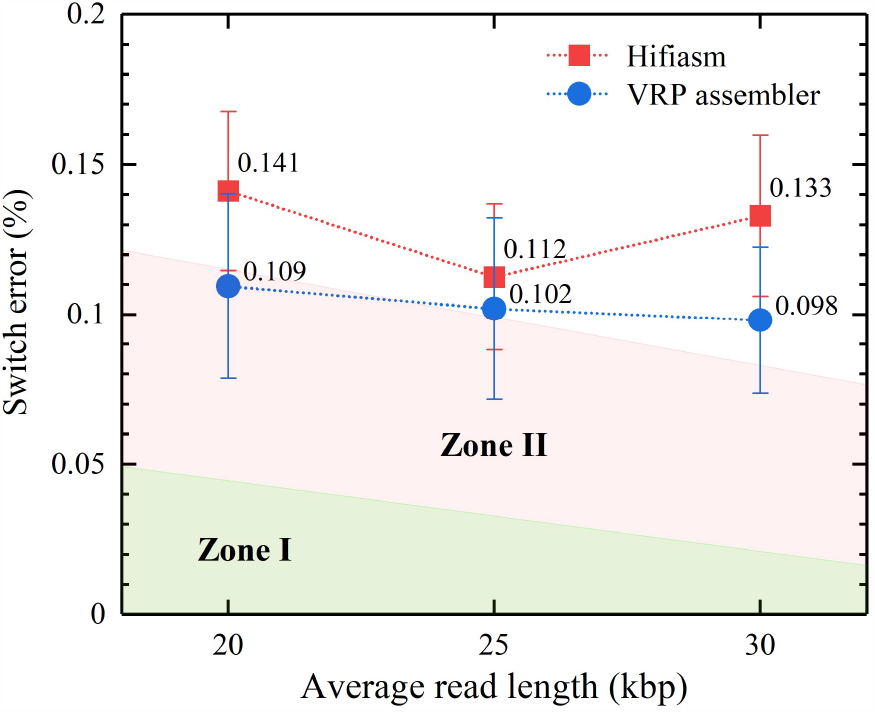
Average switch error rate of both hifiasm’s and VRP assembler’s results across three sample sets. The results of VRP assemblers is marginally better than that of hifiasm yet close to the reachable limit. Zone I is obtained by considering possible switch errors brought by distanced heterozygous SNP sites, given the N50 is manually set to be the length of MHC region. Zone II considers inevitable switch errors caused by sequencing errors. The upper bound of Zone I will approach to zero as the read length increases, while the breadth of Zone II will remain at the level of 0.06% ∼ 0.07%. In certain samples, hifiasm failed to generate accurate and valid assembly results, manifested by the assembly of two haplotypes with a combined length exceeding the total length of the MHC region thus should be classified as invalid assemblies. Therefore, during evaluation, we filtered out these invalid assemblies to ensure data integrity and reliability. The explicit results of each sample can be found in the supplementary information.

This limit is defined by two factors: the inevitable switch errors due to sequencing errors, even with correct assembly (Zone I), and the impossibility of correct phasing at heterozygous sites where the spacing exceeds the average read length (Zone II). To evaluate these two factors, we conduct a manual assembly and phasing of the reads based on the prior knowledge of the origin and order of these reads, which is obtained from their alignment positions on the reference assemblies. We use Racon to obtain the consensus sequences and evaluate switch errors hereby. Despite the theoretically perfect assembly, the results show non-zero switch errors.

This observation helps us to delineate a accuracy limit for switch errors. If the VRP assembler’s results yield a switch error rate that higher than this limit, we can confidently infer the presence of inaccurate assembly and phasing that can be improved. However, for switch error rate below this limit, one lack the necessary evidence to attribute them to either the aforementioned factors or inaccurate assembly. Hereby we delineate Zone II, which provides a valuable benchmark for evaluating the performance of our VRP assembler.

As previously discussed, factors such as sequencing errors and heterozygosity may impact the quality of assemblies. Therefore, when working with different sequencing data, it is common to choose a suitable assembler with algorithms and parameters tailored to the specific data source. For example, while hifiasm is an ideal choice for assembling PacBio CCS (HiFi) reads, its core strategy was not initially designed for handling noisy data [15, 18] or low depth of reads as in our simulated samples. Similarly, we anticipate that the VRP assembler will fully demonstrate its capabilities with even more advanced sequencing technologies that produce more accurate and longer reads.

## 3. Discussion

Although the aforementioned experiments are mainly focus on diploid genome, the VRP assembler may also conduct haplotype assembly on polyploid genomes. The ploidy of genome is essentially the number of vehicles in our model (Methods), and the VRP assembler should be able to solve triploid assembly and beyond if it could cope with diploid assembly. However, the presented assembler is currently in a stage of “toy model” and has several key points to consider.

### Intrinsic problem complexity

Given the underlying VRP model, the complexity of VRP assembler also scales as 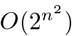, with the number of binary variables being of order *O*(*n*^2^), making it computationally expensive to locate the global optimal at large problem size. Besides, a suboptimal solution may leads to flaws such as switch errors or incorrect assembly order. As a result, obtaining the global optimal is the key in achieving the exactly correct assembly result, which is still challenging for current optimizers, including classical optimization solvers and noisy quantum devices.

### Sequencing depth

The complexity of VRP assembler grows exponentially with the number of reads, which currently limits its capability for large-scale problems on noisy intermediate-scale quantum processors due to a lack of qubits and sparse couplings. Therefore, if one aims to reconstruct longer sequences, the assembler may only deal with a lower depth of sequencing to maintain a small number of reads. Additionally, the VRP assembler is sensitive to the deviation of reads length and overlap length between reads. High deviation on reads data may invalidate the scoring function. Thus, it is advisable to filter the ideal low-depth reads from high-depth data as this approach provides a higher chance to filter out data with low deviation on reads length overlaps, which can aid in correct assembly.

### Heterozygosity rate

The VRP assembler may be more suitable for SNP-enriched genome. Like other haplotype assemblers, it needs heterozygous SNP sites in the overlap regions to distinguish the correct linkages between reads. A low rate of heterozygosity may lead to a number of switch errors and incorrect reads linkages. However, this problem can be alleviated by long and accurate sequencing technology that covers as many heterozygous SNP as possible on each reads and bring fewer sequencing errors. Additionally, the underlying optimization model of the VRP assembler may alter to adapt extra information and build accurate linkage of reads.

### Extra information

As previously discussed, switch errors are likely to occur at SNP-sparse areas. In such cases, the assembler can not determine the optimal read to link, which can hinder accurate phasing and assembly results. However, the advent of advanced sequencing technology that produces longer and more accurate reads may mitigate this issue. In deeper aspect, to further improve phasing and assembly results across SNP-sparse regions, additional information is needed to connect sparse heterozygous sites. Therefore, utilizing Hi-C data to bridge such sites may be a viable solution.

In mathematical terms, the VRP with pickups and deliveries (VRPPD) can be used to model the Hi-C integrated haplotype assembly, and potentially produce better haplotype assembly outcomes (Methods). The process can be visualized as in Fig. 6. To demonstrate the effectiveness of VRPPD model in correcting switch errors, we conduct a test to correct the reads switching in experiments on human MHC region. We choose to re-run the sample 3 with simulated reads length 30 kbp (see Supplementary Table 1). The original result contains multiple blocks of long switch between two haplotypes. Correcting these switches would enhance the accuracy of phasing. Therefore, we introduce the “pickups and deliveries” read pairs, with the expectation that reads in each pair should be phased in the same haplotype. The final assembly result demonstrates that the VRPPD model successfully corrects the mentioned long switches and reduces the switch errors from 0.13579% to 0.1164%. Therefore it confirms the efficacy of VRPPD model and its ability to enhance the phasing accuracy in haplotype assembly. The explicit results can be found in the supplementary information.

**Fig. 6.**
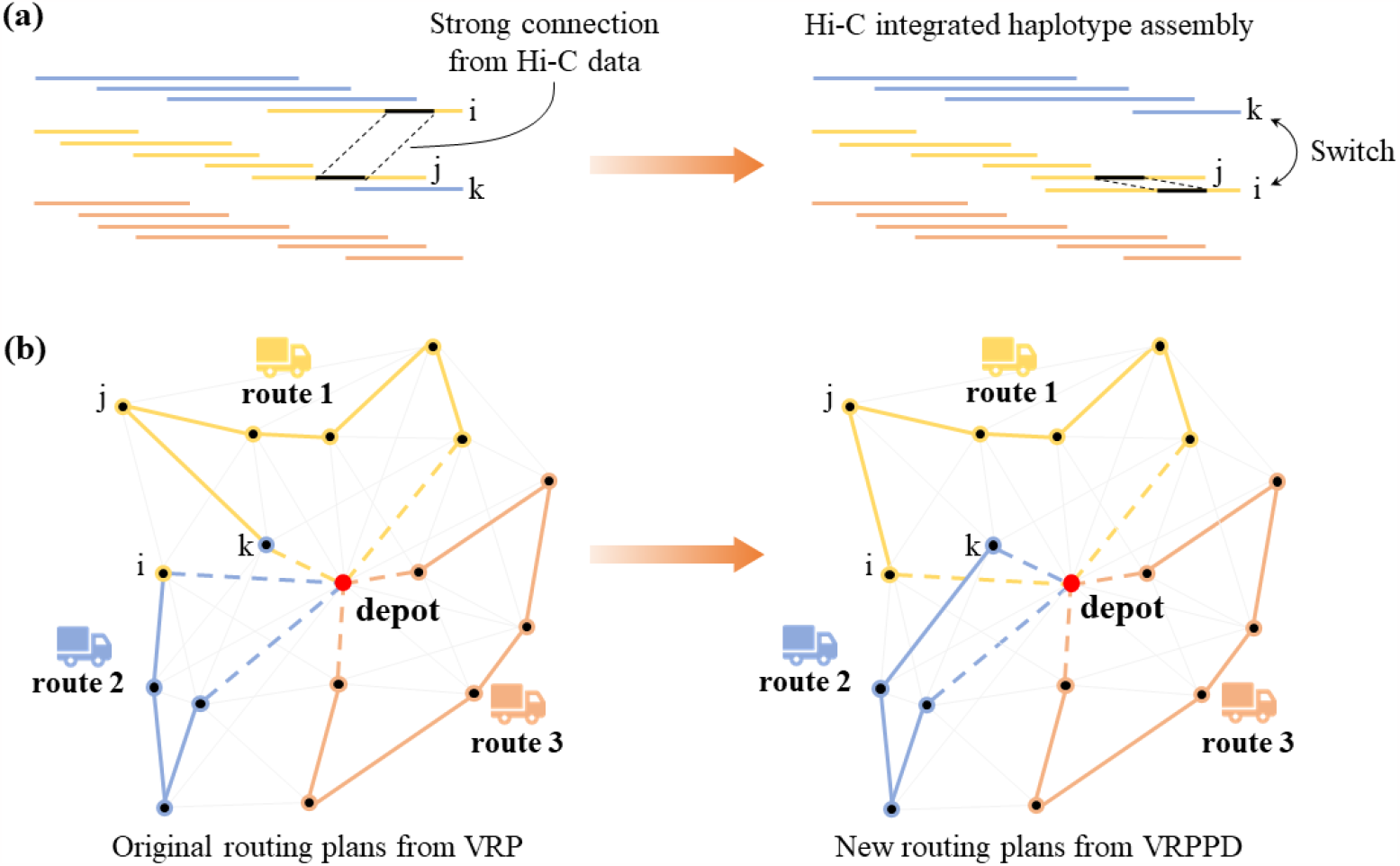
Hi-C integrated VRP assembler using VRPPD model. (a) When a Hi-C data indicates a connection between read *i* and read *j*, it suggests that these two reads may originate from the same haplotype. Given this information, VRP assembler may come out with a new assembly as shown in the right panel, where read *i* is switched with read *k* and follows read *j*. This switch can improve the accuracy of haplotype assembly by enabling the assembler to more effectively bridge sparse heterozygous sites. (b) Correspondingly in VRP model, a similar restriction limited customer *j* to be visited by the same vehicle as customer *i*. By enforcing this constraint, the optimal solution may change, resulting in customer *k* being assigned to route 2 as we expect.

Taking into account these limitations and areas for improvement, it seems VRP assembler may not yet be ready for large-scale assembly tasks using existing computational resources. Nevertheless, we look forward to the acceleration brought by mature quantum technology. Long coherence time, large qubit system as well as high fidelity qubit control and measurement are expected to significantly boost the ability of finding the global optima in complex combinatorial problems. Even if the result does not 100% guarantee the global optima, it still trends to out-perform heuristic and metaheuristic approaches in both computation time and solution quality, given the annealing time on a D-Wave’s quantum processing unit scales to nanoseconds [23]. This may enable us to design a quantum-classical hybrid strategy to further approach the global optima based on the outcomes from quantum computers.

## 4. Conclusion

In this study, we introduce the VRP assembler, a new haplotype-resolved assembly formulation based on the VRP model that can be implemented on both quantum annealer and classical optimizer.

To test the efficacy of the quantum application, we first run the VRP assembler on the D-Wave quantum annealer to perform haplotype assembly on synthetic diploid and triploid genomes. Sequencing errors are simulated to test the tolerance of the assembler. Additionally, we test the VRP assembler with Google’s OR-Tools for reconstructing a real human genome segment, the MHC region, to evaluate its performance under the realistic heterozygosity rate and the degree of genome duplication.

Our results show that the VRP assembler can solve haplotype assembly problem of small size genome on the current available quantum annealer. We analyze that VRP assembler may perform better under specific conditions, such as high-accuracy reads with similar lengths from advanced sequencing technology and high heterozygosity rate from appropriate target genome. Furthermore, we note that sparse heterozygous SNP sites may limit the VRP assembler’s ability to produce proper results. As a potential solution, we suggest using the VRPPD model to reduce switch errors by integrating Hi-C data, although this approach may increase computational costs. We also plan to explore the VRPPD model’s capabilities in the future studies.

In conclusion, the VRP assembler presents a potential alternatives for haplotype-resolve assembly despite several limitations. Since the limitations mainly arise from the computational complexity of VRP model, we anticipate that more advanced quantum technology will provide better solutions in the future.

## 5. Methods

In this section, we illustrate the details of the VRP assembler, including formulations of VRP and procedures of mapping a haplotype assembly problem into a VRP. Moreover, we extract general background of quantum optimization and the unconstrained model of VRP that can be processed on a quantum annealer.

### 5.1 Vehicle routing problem

In combinatorial optimization, VRP is a mix integer linear programming (MILP) problem [48]. It models a real world scenario where a fleet of vehicles try to find the optimal set of routes to deliver goods to a number of customers. Additional constraints can be integrated into the model to meet a more practical need, such as capacity of vehicles, time windows of customers, multiple starting and ending point, etc. Here we only utilize the simplest case without the conditions above. The formulation is

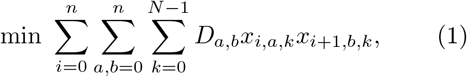

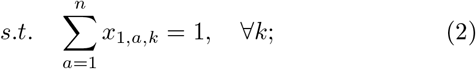

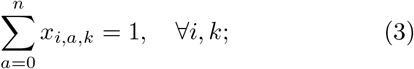

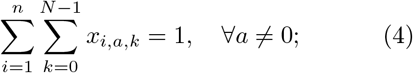

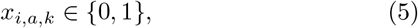

where *N* is the number of vehicle routes, *n* is the number of customers (or nodes). The set of *x*_*i,a,k*_ represents decision variables, where 0 ≤ *i* ≤ *n* + 1, 0 ≤*a* ≤*n* and 0 ≤*k* ≤*N* −1. They encodes whether vehicle *k* visits the customer *a* at its *i*-th step. If true then *x*_*i,a,k*_ = 1, otherwise *x*_*i,a,k*_ = 0. Equations (2) to (4) are the constraints. Note that the 0-th step and (*n* + 1)-th step should be initialized on the depot (*a* = 0) by setting *x*_0,0,*k*_ = *x*_*n*+1,0,*k*_ = 1 for all *k*, and every vehicle is ensured to leave the depot at their first step by Eq. (2). Equation (3) makes sure that each vehicle can only be at one place at each step. Equation (4) constrains that every customer will be visited once and only once. The depot can be visited multiple times by multiple vehicles. Equation (1) is the objective function of total travelling distance of *N* vehicle routes, which we aim to minimize under the above constraints.

#### 5.1.1 Vehicle routing problem with pickups and deliveries

In VRP with pickups and deliveries (VRPPD), vehicles pick up goods from one customer and deliver them to another customer in the middle of their routing plan. Both customer from such a customer pair must meet two conditions: First, they must be visited by the same vehicle. Next, the customer at the pickup spot must be visited ahead of the delivery spot. However, since we can not infer the relative order of two reads in any way before conducting assembly, we may omit the second condition and concentrate on the first one. In mathematics, we may add the following expression to the original formulation of VRP Equations (1) to (5),

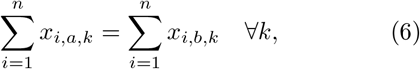

where *a* and *b* are from the pickup-delivery customer pairs set.

### 5.2 Mapping haplotype assembly to VRP

The optimization of VRP is to find the minimum total distance of *N* routes. All routes diverge from the same depot, get all customers visited once and then eventually converge at the same depot. To translate the haplotype assembly problem to a VRP, we outline the following four steps:

i. Set the number of routes *N* equal to the ploidy of the genome being studied;
ii. Set the number of customers *n* to be the number of sequencing reads;
iii. Determine weighted edges by a scoring function as

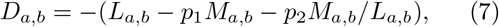

where *L*_*a,b*_ corresponds to the *alignment block length* between read *a* and read *b, M*_*a,b*_ represents the *number of mismatches* within the alignment block; *p*_1_ and *p*_2_ are the penalty parameters for reducing scores associated with mismatches. The needed *alignment block length* and the *number of mismatches* can be obtained by pairwise alignments software, such as minimap2 [57];
iv. The depot is considered as a virtual read and has the same score aligned with all real reads, that is

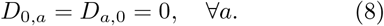

The first two steps lead that the customers visited by the same vehicle corresponds to reads sorted in the same haplotype. Moreover, the visiting order of customers indicates the assembly order of reads. As previously mentioned, we employed SimLoRD to generate low-depth reads for each sample. Due to computational resource limitations, VRP assembler can currently only handle low-depth reads. However, directly generating low-depth of reads from the reference haplotypes using SimLoRD often results in incomplete coverage of the MHC region, rendering VRP assembler ineffective. Therefore, we initially generate 50X reads using SimLoRD, which gives full coverage of the MHC region. Subsequently, we filtered out 4 ∼ 6X reads that still provided complete coverage of the MHC region.

To accomplish this, we first align all reads to the reference haplotypes. Then, based on the length of the reads, their alignment positions, and alignment qualities, we iteratively selected suitable reads. The same procedure was applied to both reference haplotypes. Finally, the generated read files from the two haplotypes were merged, resulting in a low-depth reads file that ensured complete coverage of the MHC region. This reads generation process was repeated for each sample, and the identical reads file was then provided as input to both VRP assembler and hifiasm. Step (iii) is essential to realize phasing and assembly simultaneously. Step (iv) turns the cycle problem in original VRP into the path problem that consists with *de novo* assembly. These four steps build up the procedure from Figure 1(a) to Figure 1(b).

The details in step (iii) are worth mentioning. From the BAM file generated by minimap2, tag “NM” tells the *number of mismatches* inside the alignment block between reads. These mismatches may arise due to sequencing errors, heterozygosity, INDEL, and most commonly reads originating from distinct position. On one hand, heterozygosity and INDEL would result in greater number of mismatches between reads from different alleles. As such, these genetic variations may be leveraged for phasing purposes. On the other hand, the number of mismatches is obviously lower between reads around the same position, thus can be used in assembly. However, not all information from the original BAM file are valuable. There may be many undesirable alignments in the output BAM file, such as multiple alignment positions between two reads, and the redundant information between reads from distinct position. Besides, eliminating redundant alignments may result in a sparser VRP graph, which is easier to solve. Therefore, a filtering strategy is implemented to screen out a clean alignment information as illustrated in Figure 7. The strategy includes three steps as follows:

**Fig. 7.**
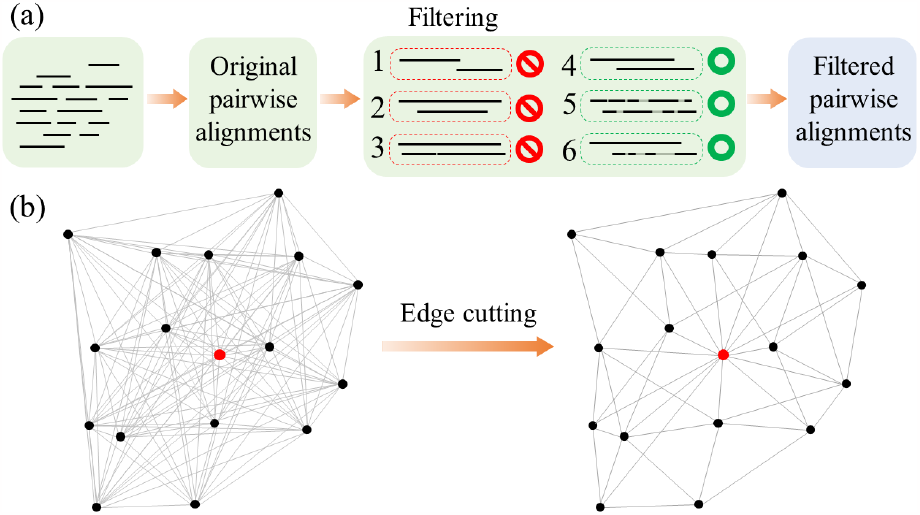
The filtering strategy of pairwise alignment file. (a) We use minimap2 to obtain an pairwise alignment file. Then we filter out three types of unhelpful alignment: 1. too short alignment block, 2. the alignment block being longer or equal to the reads, and 3. the alignment block being shorter but very close to the reads length. Additionally, there may be multiple alignment positions within the same reads pair, but only one with highest alignment quality should be kept. After that, the simplified alignment information are used as the input of the VRP assembler. (b) Essentially, the entire filtering process is to cut particular edges in a dense graph, making it easier to solve.

i. Alignments with too short alignment block, which indicates that the query read and the target read are not adjacent to each other, are moved.
ii. Alignments with an alignment block that is longer or equal to one of the reads are filtered out as this information is redundant.
iii. Alignments with a too long alignment block that is close to the reads itself are eliminated. Although these may offer minor help in extending sequence, the aligned reads are likely to come from a similar position of different haplotypes, and hence, they should not be linked.

#### 5.2.1 Integrating Hi-C data with VRPPD

As mentioned in Section 3, Hi-C information can be integrated to help enhancing accuracy of haplotype assembly. However, a typical line from Hi-C data may only suggest that the two involved DNA segments are likely to originate from the same haplotype, without providing information on their relative positions. Therefore, incorporating Hi-C data in VRP assembler requires only the first condition listed in Section 5.1.1, or Equation (6) where *a* and *b* represent read *a* and read *b* originate from the two ends of a Hi-C data.

### 5.3 Quantum annealing for VRP

The search space of VRP is at least 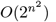, making it computationally intractable to locate the global optimum in a large problem scale. Heuristic methods such as cheapest insertion and savings algorithm, are generally fast but most likely to be trapped in a local minima. To cope with this situation, metaheuristic approaches such as simulated annealing and guided local search, although do not guarantee the optimality, offer acceptable balance between the solution quality and computational time by escaping local minimum rather efficiently.

There are various strategies to further enhance the optimization performance. For example, a feasible result from a heuristic method can in turn serve as a starting point in a subsequent metaheuristic procedure, thereby reducing the time required to reach a good solution. Nonetheless, in large and complex instances, these algorithms almost inevitably fall in suboptimal situations and can hardly converge into better states.

Quantum adiabatic computation shows its potential to help get out of the dilemma. The speedup of quantum annealing over metaheuristic methods has been revealed on particular tasks [58–60]. Optimizations can be solved by performing an adiabatic evolution of an corresponding Ising model on a quantum annealer like D-Wave’s system. The evolution of the core system in quantum annealers may be described by a time-dependent Hamiltonian of transverse-field Ising model,

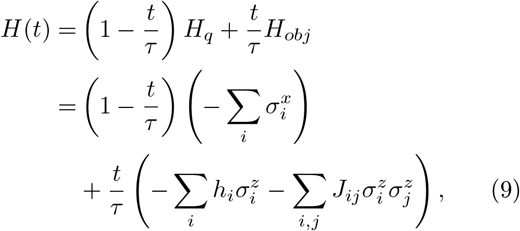

where *h*_*i*_ and *J*_*ij*_ are the local fields and interaction strengths on qubits, respectively. 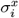 and 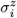are Pauli-X and Pauli-Z operators on qubit *i. H*_*q*_ term provides an external transverse field that induces quantum fluctuation. *H*_*obj*_ encodes the target problem to be solved. *τ* is the annealing time and should be set with respect to the energy gap between the ground state and the first excited state [61].

As per the quantum adiabatic theorem [61, 62], if a quantum system is initially in its ground state of *H*(0) and if *H*(*t*) changes slowly enough during the adiabatic process, the system *H*(*t*) will remain in its ground state throughout the evolution, despite changes in the energy landscape as shown in Fig. 8. This is owing to the effects of quantum fluctuations and quantum tunneling, which enable the system to traverse the energy barrier and stay in its ground state. More formally, with increasing *τ*, the probability of finding the system *H*(*τ*) in its ground state at the end of evolution will approximate to unity as ∼ 1 −*τ* ^−2^ [61].

**Fig. 8.**
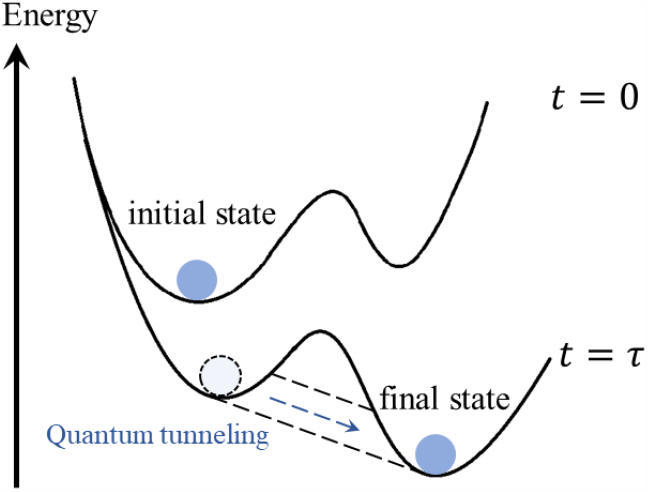
Quantum adiabatic evolution in quantum annealing. During an evolution, the energy landscape of the quantum system may alter, leading to modifications of the system’s ground state. However, if the evolution time *τ* is sufficiently long, the Hamiltonian that governs the system will vary slowly, allowing the system to search for and transfer to a new ground state via the quantum tunneling effect.

Ultimately, the system will converge towards the global minimum of *H*(*τ*) = *H*_*obj*_ with energy

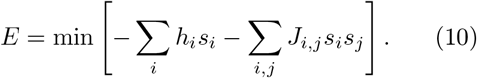

Here, *s*_*i*_ ∈{+1, −1} is the eigenvalue of Pauli-Z operator on qubit *i*.

Therefore, quantum annealing is a natural approach for solving combinatorial optimization problems, provided that one can map the target problem to *H*_*obj*_. As a result, in order to solve the target problem on a quantum annealer, one must transform the problem into formulation as Eq. (10), or equivalently, the quadratic unconstrained binary optimization (QUBO) form as

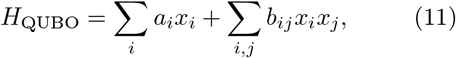

where *x*_*i*_ ∈{0, 1}. *a*_*i*_ and *b*_*ij*_ are bias and coupling, respectively.

Many optimization problems can be formulated into QUBO in multiple ways [63]. Here, we use the developed framework in [64] and formulate VRP in Equations (1)-(5) into QUBO as follows

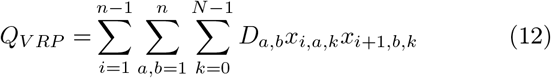

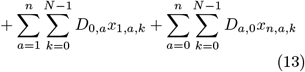

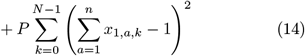

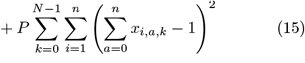

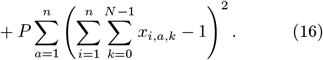

Objective term (12) and (13) are of the same form as the objective function in Eq. (1). Note that *x*_0,0,*k*_ and *x*_*n*+1,0,*k*_ are both fixed to 1, thus all terms involving *x*_0,*a,k*_ and *x*_*n*+1,*a,k*_ can be omitted. Besides, term (13) offers zero contribution according to the settings in Equation (8). Constraints in Eq. (2) are represented by terms (14). The penalty terms (15) and (16) encode constraints Eqs. (3) and (4), with the penalty parameter *P* being set to 10^7^.

The above QUBO formulation will be transformed into Ising model as Equation (9) by the D-Wave’s Ocean software automatically and submitted to its quantum processing unit (QPU) on cloud. The returning configuration will be decoded via the framework in [64] and will be further translated into the assembly results in different haplotypes as illustrated in Figs. 1(c) and 1(d).

To sum up, a haplotype assembly problem can be solved on a quantum annealer by mainly five steps:(i) generate a pairwise alignment file from sequencing reads; (ii) filter out redundant alignments; (iii) build a VRP instance in QUBO formulation; (iv) upload the problem onto the D-Wave’s quantum annealing system; (v) decode the returning configurations.

## Data availability

All relevant data supporting this study are available within the article and its supplementary material or from the corresponding author upon reasonable request. The assembled reference haplotigs reported in Ref.[2] can be obtained from https://github.com/NCBI-Hackathons/TheHumanPangenome/tree/master/MHC/assembly/MHCv1.1. The simulated reads and the corresponding assemblies are available at https://github.com/yiboch/vrp_assembler/tree/main/data.

## Code availability

Scripts of the filtering strategy, the switch error evaluation and the commands for necessary softwares are available at https://github.com/yiboch/vrp_assembler/tree/main/scripts. All codes supporting this study are available upon reasonable request.

## Supplementary information

The supplementary information is enclosed at the end of this file.

## Acknowledgments

The authors would like to thank Chentao Yang, Jing-Kai Fang and Mengyang Xu for helpful discussions. This work is supported by the National Key R&D Program of China (2022YFC3400400). This work is also supported by the China National GeneBank, and the Guangdong Bigdata Engineering Technology Research Center for Life Sciences.

## SUPPLEMENTARY INFORMATION

### 1 Switch error rate evaluation

**Table 1:**
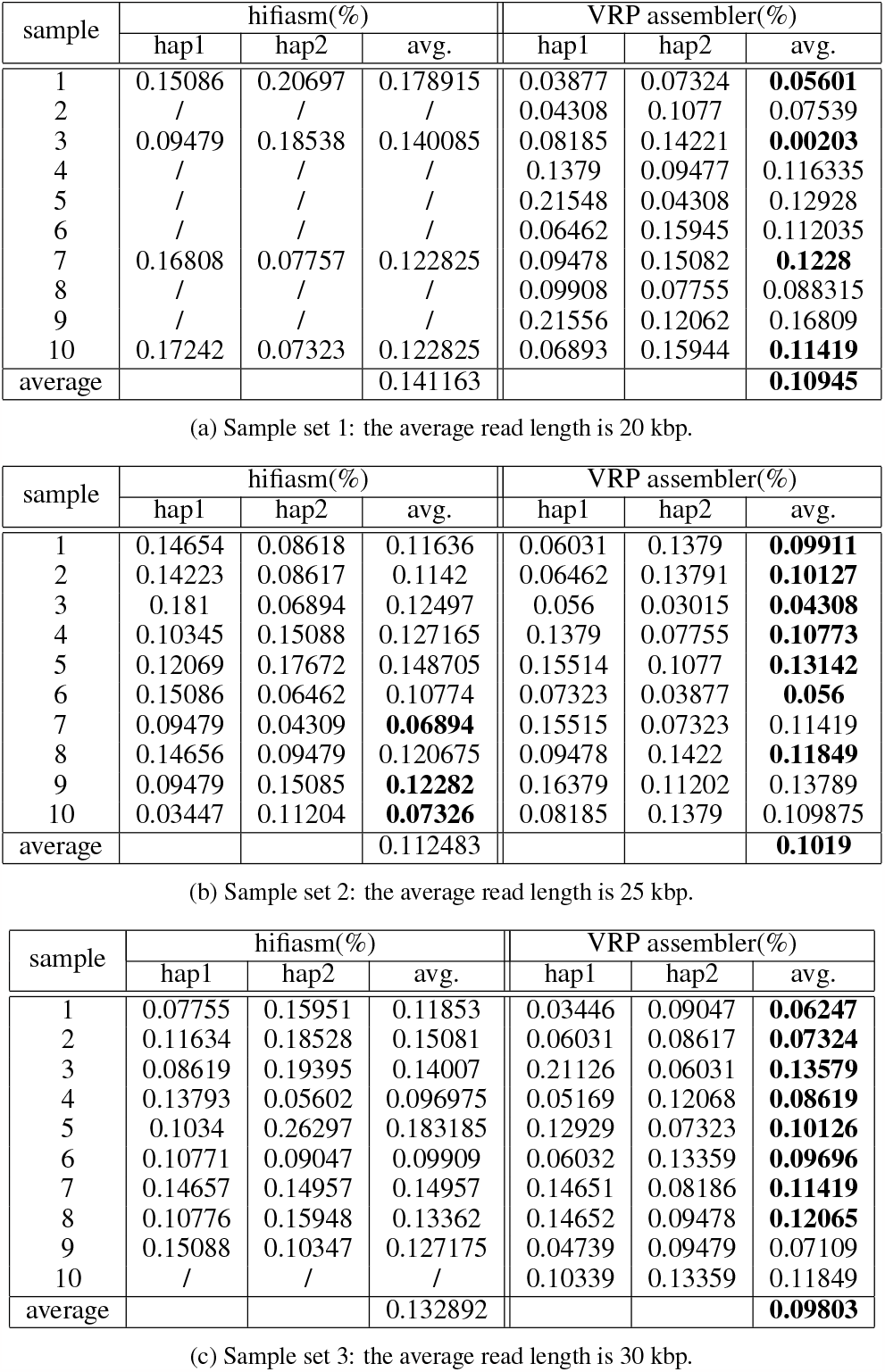
Switch error rate comparison of hifiasm and VRP assembler on three sample sets with different average read length. Data with “/” means incorrect assembly with total assembled length greater than the entire MHC region (∼5Mbp). Bold text represents a better performance.

### 2 Proof of concept of VRPPD

For the third sample in the 30kbp sample set, the original reads assembly order obtained by ortools is shown in Figure 1, reads index with yellow marked are believed to be phased incorrectly, which would bring switch errors in the final assembly sequence.

**Figure 1.**
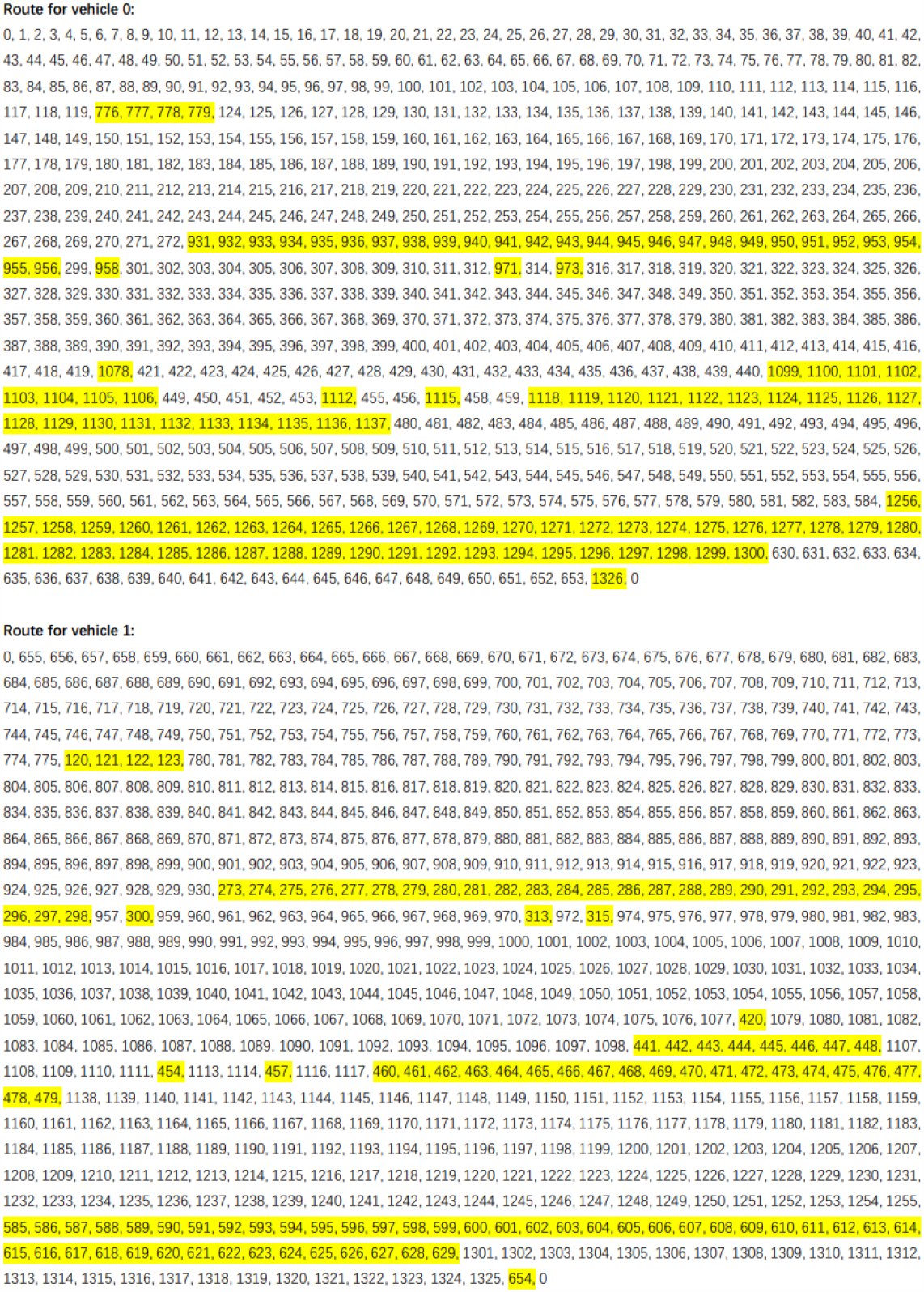
Original outputs of the third sample in sample set 3 without pickups and deliveries.

After we add the following pickups and deliveries pairs:

[(1255, 1256), (584, 585), (629, 630), (1300, 1301), (272, 273), (930, 931), (956, 957), (298, 299), (459, 460), (1117, 1118), (1137, 1138), (479, 480), (440, 441), (1098, 1099), (1106, 1107), (448, 449), (119, 120), (775, 776), (123, 124), (779, 780)], the new reads assembly order comes out as in Figure 2. The result shows a noticeable decrease of reads switching, suppressing the average switching error rate from 0.13579% to 0.1164%.

**Figure 2.**
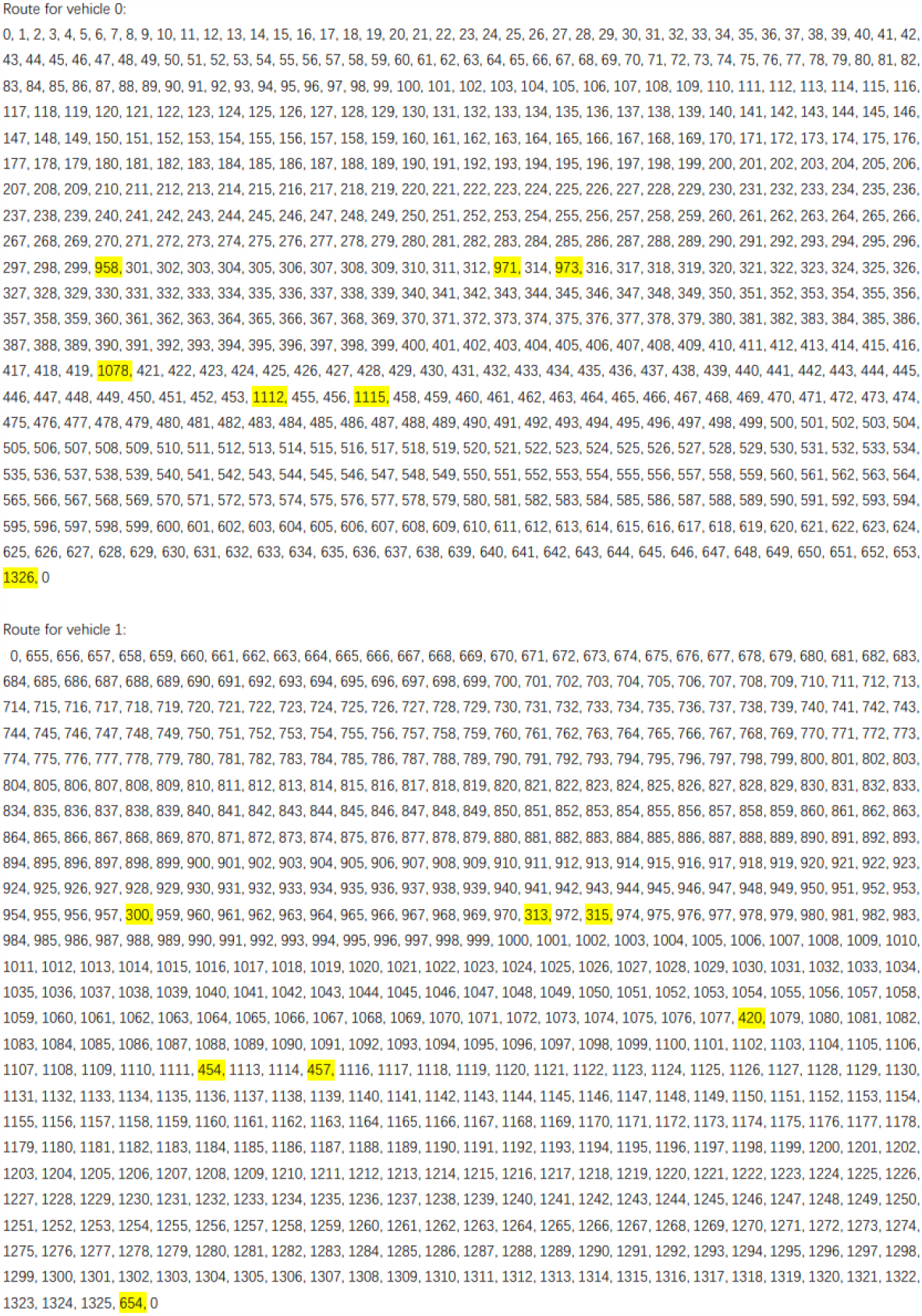
VRPPD outputs of the third sample in sample set 3 with pickups and deliveries pairs.

### 3 Scripts and parameter setting

#### 3.1 Scripts in processing relative files

The necessary commands as well as switch error scripts can be found in our code repository.

#### 3.2 Parameters in BAM filtering and alignment scoring

Section 3.2 lists all parameters when filtering BAM file and scoring alignments for each sample. The *p*_1_ and *p*_2_ are referencing to the penalty parameters in the scoring function:

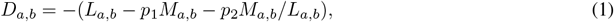

**Table 2:**
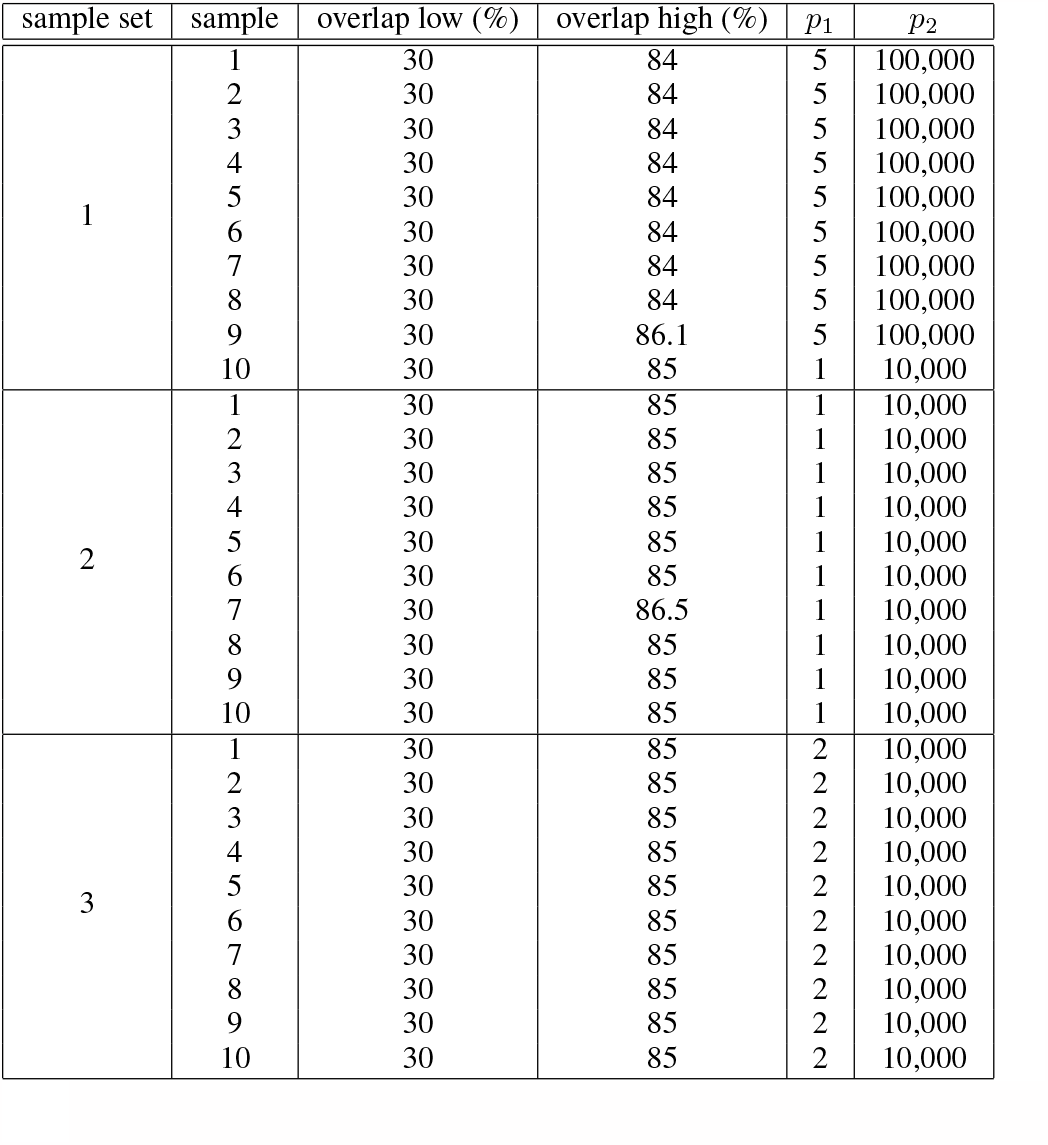
Parameters setting in scoring alignments of each sample.

## 4. Source data

All source data and all simulated reads samples can be found in our code repository.

